# Autoregulation of the *S. mutans* SloR metalloregulator is constitutive and driven by an independent promoter

**DOI:** 10.1101/301416

**Authors:** Patrick Monette, Richard Brach, Annie Cowan, Roger Winters, Jazz Weisman, Foster Seybert, Kelsey Goguen, James Chen, Arthur Glasfeld, Grace Spatafora

## Abstract

*Streptococcus mutans*, one of ∼600 bacterial species in the human oral cavity, is among the most acidogenic constituents of the plaque biofilm. Considered to be the primary causative agent of dental caries, *S. mutans* harbors a 25kDa SloR metalloregulatory protein which controls metal ion transport across the bacterial cell membrane to maintain essential metal ion homeostasis. The expression of SloR derives, in part, from transcriptional readthrough of the *sloABC* operon which encodes a Mn^2+^/Fe^2+^ ABC transport system. Herein, we describe the details of the *sloABC* promoter that drives this transcription, as well as a novel independent promoter in an intergenic region (IGR) that contributes to downstream *sloR* expression. RT-PCR studies support *sloR* transcription that is independent of *sloABC* expression, and the results of 5′ RACE revealed a *sloR* transcription start site in the IGR from which the −10 and −35 promoter regions were predicted. The results of gel mobility shift assays support direct SloR binding to the IGR, albeit with lower affinity than SloR binding to the *sloABCR* promoter. Function of the *sloR* promoter was validated in qRT-PCR experiments. Interestingly, *sloR* expression was not significantly impacted when grown in the presence of high manganese, whereas expression of the *sloABC* operon was repressed under these conditions. The results of *in vitro* transcription studies support SloR-mediated transcriptional-activation of *sloR* and -repression of *sloABC.* Taken together, these findings implicate SloR as a bifunctional regulator that represses *sloABC* promoter activity and encourages *sloR* transcription from an independent promoter.

**Importance:** Tooth decay is a ubiquitous infectious disease that is especially pervasive in underserved communities worldwide. *S. mutans*-induced carious lesions cause functional, physical, and/or aesthetic impairment in the vast majority of adults, and in 60-90% of schoolchildren in industrialized countries. Billions of dollars are spent annually on caries treatment, and productivity losses due to absenteeism from the workplace are significant. Research aimed at alleviating *S. mutans*-induced tooth decay is important because it can address the socioeconomic disparity that is associated with dental cavities and improve overall general health which is inextricably linked to oral health. Research focused on the *S. mutans* SloR metalloregulatory protein can guide the development of novel therapeutics and so alleviate the burden of dental cavities.

## Introduction

Dental caries are important indicators of oral health and overall general health in both children and adults. Despite significant public health efforts aimed at reducing caries incidence, approximately 60-90% of school-age children worldwide experience caries, with 91% of adult caries in the United States involving the permanent dentition (1, 2). Of particular concern are children of socioeconomically disadvantaged families who are twice as likely to experience rampant caries in comparison with their wealthier counterparts, and who often present with severe clinical outcomes later in life (3).

Among the early colonizers of the tooth surface is *Streptococcus mutans*, considered to be the primary causative agent of dental cavities in humans (4). Ongoing research aimed at alleviating or eliminating caries has given rise to valuable insights of *S. mutans* virulence properties, including genes that mediate its obligate biofilm lifestyle, its ability to tolerate acid and oxidative stress, and maintain metal ion homeostasis (5–12). The introduction of sucrose into the Western diet marked a turning point for *S. mutans*, which metabolizes carbohydrates for energy production and generates a lactic acid byproduct that demineralizes tooth enamel and drives the process of tooth decay (13, 14). Taken together, *S. mutans*’ arsenal of virulence attributes makes it an especially resolute dental pathogen, and in dysbiotic plaque a primary instigator of caries formation (15, 16).

Among the evolutionary responses that are paramount for *S. mutans* survival and pathogenesis in dental plaque is tight regulation of essential metal ion transport across the bacterial cell membrane. To this end, *S. mutans* is endowed with metal ion uptake machinery which enables the import of divalent cations such as Mn^2+^ and Fe^2+^ that are essential for cellular and subcellular functions, and ultimately for bacterial cell viability. Aberrant metal ion uptake, however, can result in over-accumulation of intracellular metal ions and bacterial cell death owing to Fenton chemistry and the elaboration of toxic oxygen radicals (17–20). To counteract these deleterious effects and achieve intracellular homeostasis, *S. mutans* has evolved transport mechanisms that function to maintain appropriate metal ion uptake, which is especially paramount during periods of feast and famine in the oral cavity when Mn^2+^ concentrations can vary greatly.

*S. mutans* metal ion uptake is mediated, in large part, by the *sloABCR* operon which encodes a SloABC Mn^2+^/Fe^2+^ transporter and, via transcriptional readthrough, a 25 kDa SloR metalloregulatory protein. As a transcription factor, SloR, modulates metal ion transport upon binding to DNA in response to manganese availability (7, 20, 21). For instance, between mealtimes exogenous metal ions are not readily available because they are sequestered to host proteins, such as lactoferrin in saliva. Hence, under fasting conditions, *S. mutans* upregulates the *sloABC* gene products which includes ATP-binding and -hydrolyzing proteins as well as a transmembrane SloC lipoprotein that scavenges metal ions for uptake (20, 21). In contrast, during a mealtime free metal ions are plentiful in the oral cavity, and in response *S. mutans* downregulates its metal ion importers so as to avoid over-accumulation of intracellular metal ions and their associated cytotoxic effects. We believe SloR mediates such metalloregulation by binding directly to Mn^2+^ which, in turn, promotes SloR dimerization and a subsequent conformational change at the N-terminus of the protein to facilitate DNA binding. Specifically, in a previous report we describe SloR-DNA binding upstream of the *sloABC* locus to a promoter-proximal SloR Recognition Element (SRE) that represses *sloABC* transcription, presumably via a mechanism that involves promoter exclusion to RNA polymerase (RNAP) (22, 23). Hence, the *sloABC*-encoded metal ion uptake machinery in *S. mutans* is subject to transcriptional repression by SloR under conditions of Mn^2+^ availability, and conversely to de-repression when Mn^2+^ becomes limiting. The transcriptional regulatory properties of the SloR protein are not limited to the *sloABC* locus. In fact, work in our laboratory suggests that the SloR protein, a member of the DtxR family of metalloregulators, may be involved in regulating as many as 200 genes in the *S. mutans* genome, either directly or indirectly (24). The genes that are subject to SloR control belong to a variety of different functional categories beyond metal ion homeostasis, and include gene products that mediate *S. mutans* oxidative stress and acid tolerance, biofilm formation, and genetic competence, all of which contribute to *S. mutans* virulence (7, 8, 24, 25). In addition, accumulating evidence in our laboratory supports SloR as more than just a repressor of *S. mutans* gene expression. While the results of expression profiling studies support SloR-mediated-repression of certain genes (called Class I genes), the binding of SloR to other gene loci can culminate in gene activation (called Class II genes) (24). The mechanism by which SloR encourages gene transcription is unknown and is currently under investigation in our laboratory.

Despite the central importance of SloR in promoting *S. mutans* survival and virulence gene expression, surprisingly little is known of the regulatory mechanism(s) that modulate SloR itself. To date, the regulation of SloR in *S. mutans* has been shown to be manganese-dependent and driven, in part, by the *sloABC* promoter via transcriptional read-through of a weak terminator that is located 3’ to the *sloC* gene (7, 20, 22). Hence, SloR levels that derive from *sloABC* promoter activity likely vary with Mn^2+^ availability between and during meal times. We propose however, that some constitutive baseline level of SloR is likely necessary to facilitate scavenging of essential Mn^2+^ and/or Fe^2+^ by the SloC metal ion importer regardless of exogenous metal ion availability, and that such fine-tuning could involve additional mechanisms of control.

In the present study, we set out to determine whether a 184 base pair intergenic region (IGR) that is located immediately downstream of the *sloC* coding region and upstream of the *sloR* gene, harbors a specific promoter that drives *sloR* expression independent of *sloABC* promoter activity. We propose that together with a unique and as-yet-unidentified SRE in this IGR, a bifunctional role for SloR as both a repressor and activator of *S. mutans* gene expression will be supported (26).

## Results

### SloR homodimers bind to a 72bp SRE

In a previous report, we describe the thermodynamic binding properties of the *S. mutans* SloR metalloregulator to its cognate SRE within the *sloABC* promoter region. Herein, the results of fluorescence anisotropy studies conducted with 1mM manganese and various SRE-containing DNA fragments revealed tight binding of the SloR protein to a 72bp DNA fragment (Kd = 32nM). Combined with the results of EMSA and DNA footprinting experiments which support the binding of at least two SloR dimers to this 72bp sequence, we hypothesized that SloR may bind as a set of homodimers to each of three inverted hexameric repeats on this 72bp target DNA, each with the consensus sequence AATTAA or some modification thereof (Figure 1). To test this hypothesis, we used the SloR protein, 1mM Mn^2+^, and the predicted 72bp SloR recognition element in negative staining and electron microscopy experiments. The resulting dataset comprised of 55 total images was classified into two-dimensional class averages using a PARTICLE software package (www.image-analysis.net/EM/). The dominant pattern that was revealed by the class averages share a common shape with three distinct ellipsoidal regions presumed to be SloR dimers (Figure 1 inset, arrowheads), tilted off of the DNA axis by ∼32°. The SloR binding pattern occupies a total length of 239 Angstroms on the DNA with each SloR dimer measuring ∼90 Angstroms across, and the distance from the center of one dimer to the center of the next measures 75 Angstroms (equal to 22bp). Taken together, the binding pattern and low-resolution image of the SloR-SRE interaction are consistent with the binding of three SloR dimers to the 72bp target sequence.

**Figure 1.**
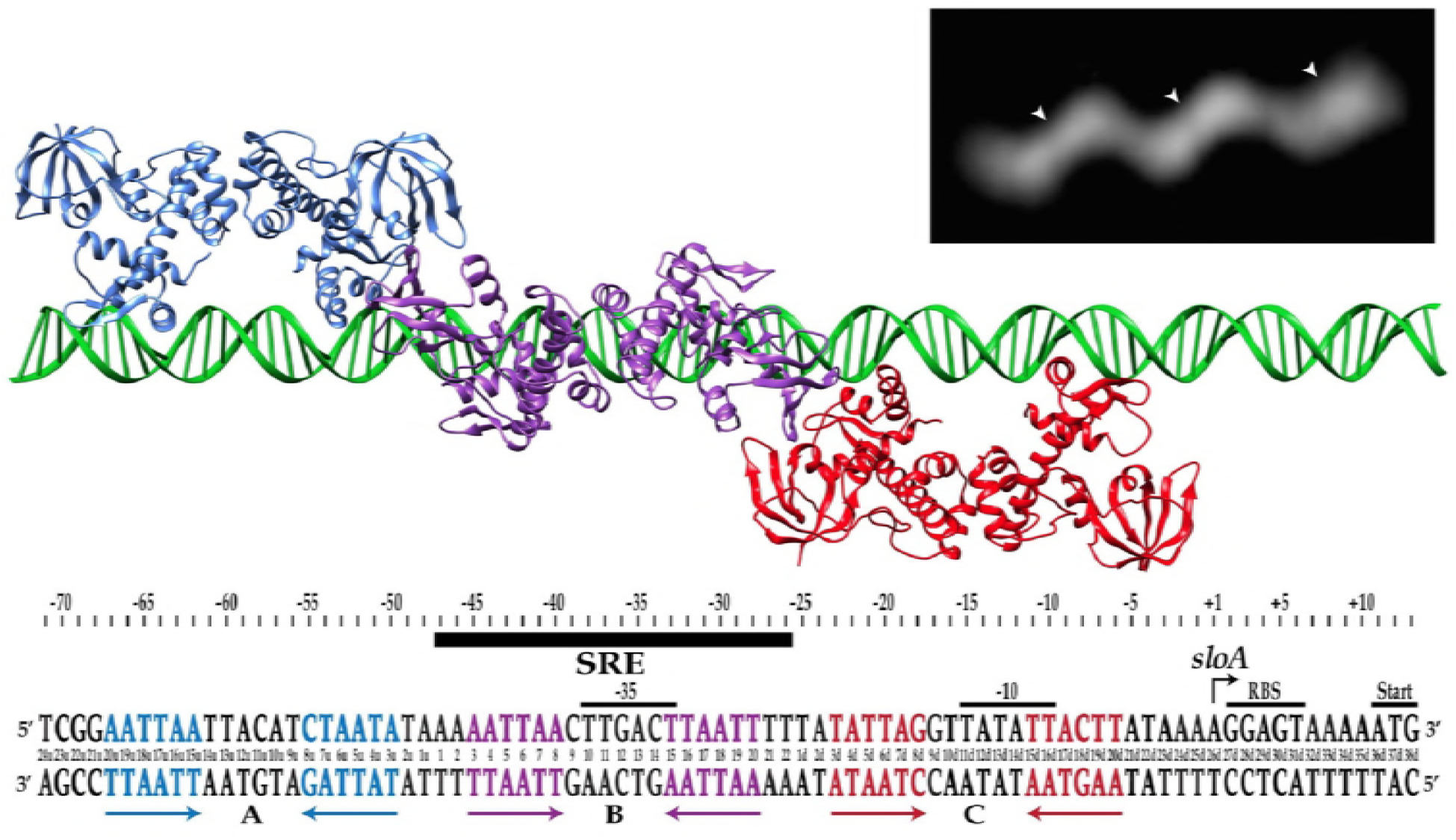
SloR-SRE binding model in the region of the *sloABC* promoter. Shown are each of three inverted hexameric repeats (in blue, purple, and red) that span 72bp in the *sloABC* promoter region, and to which SloR presumably binds as three homodimers. Affinity binding studies support preferential binding of SloR homodimers to the hexameric repeat in region B (designated by the black bar), followed by cooperative binding of SloR dimers to regions A and C. Inset: Negative staining of the SloR-SRE interaction *in vitro* supports homodimeric binding of SloR to a 72bp SloR recognition element (SRE). Shown are three SloR homodimers bound to a DNA filament containing the 72bp SloR binding element. The arrowheads denote the center of mass for each SloR homodimer.

### SloR binding to the 72bp SRE is cooperative

Equilibrium binding of SloR to fluoresceinated duplex DNA containing sequences from the *sloABC* promoter region (Table S1) was measured using fluorescence anisotropy with SloR binding to fluorescently-labeled duplex DNA containing either one, two, or three 22bp sequences identified previously on the 72bp SRE (labeled A, B and C in Figure 1). Region B, which includes a pair of inverted repeats, was the only site that demonstrated SloR-specific binding under conditions as high as 250mM NaCl (data not shown). Sequences A and C, which deviate from the consensus 22bp sequence at two and three nucleotide positions respectively, do not measurably bind SloR under these same assay conditions. Saturation binding to these sequences was observed however, when the salt concentration was lowered to 50mM, albeit along with some non-specific interactions. With that said, when SloR was titrated into solutions containing 46bp duplexes harboring two contiguous pairs of inverted repeats, either *sloA*_AB or *sloA*_BC, specific cooperative binding was observed under high salt conditions, with Hill coefficients of 1.8 and 1.7, respectively. This result indicates that SloR is capable of binding to the A and C sites with high affinity if the B site is already occupied, consistent with SloR interactions at the B-site that recruit the additional dimers to the flanking A and C sites. In addition, we measured 50% occupancy of the two sites on *sloA*_AB and *sloA*_BC at 26nM and 10nM SloR, respectively (data not shown). This result corroborates the observed binding of SloR dimers to all three sites on the 72bp duplex described above in the negative staining experiments.

### Differential expression of the *S. mutans sloABC* and *sloR* genes suggests the presence of a *sloR*-specific promoter

To determine whether expression of the *sloABC* and *sloR* genes is coordinated under conditions of low versus high manganese, we performed qRT-PCR experiments, the results of which reveal different transcription profiles (Table 2) despite the derivation of these genes from a single polycistronic mRNA. Specifically, expression of the *sloABC* operon was repressed 3-fold under conditions of high manganese availability (Student’s *t*-test, p<0.05), whereas transcription of the *sloR* gene was not significantly altered under these same conditions (Student’s *t*-test, p>0.05). That expression of the *sloABC* and *sloR* genes is different under conditions of high Mn^2+^ suggests an additional control mechanism for *sloR* transcription that may involve an independent promoter. We predict this promoter is located within the 184bp intergenic region (IGR) that separates the *sloC* gene of the *sloABC* operon from the downstream *sloR* gene on the *S. mutans* chromosome.

**Table 1.**
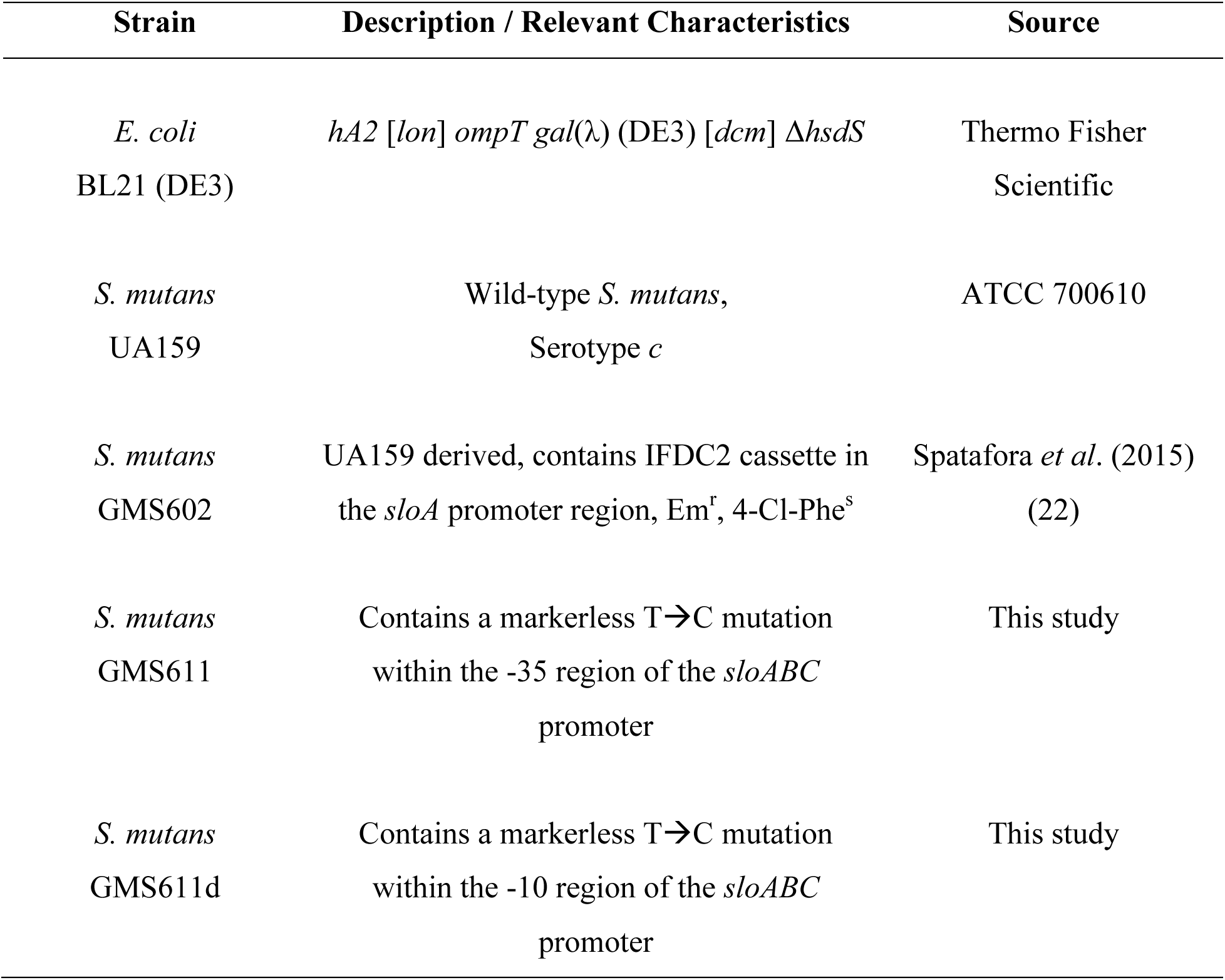
Bacterial strains used in this study.

**Table 2.**
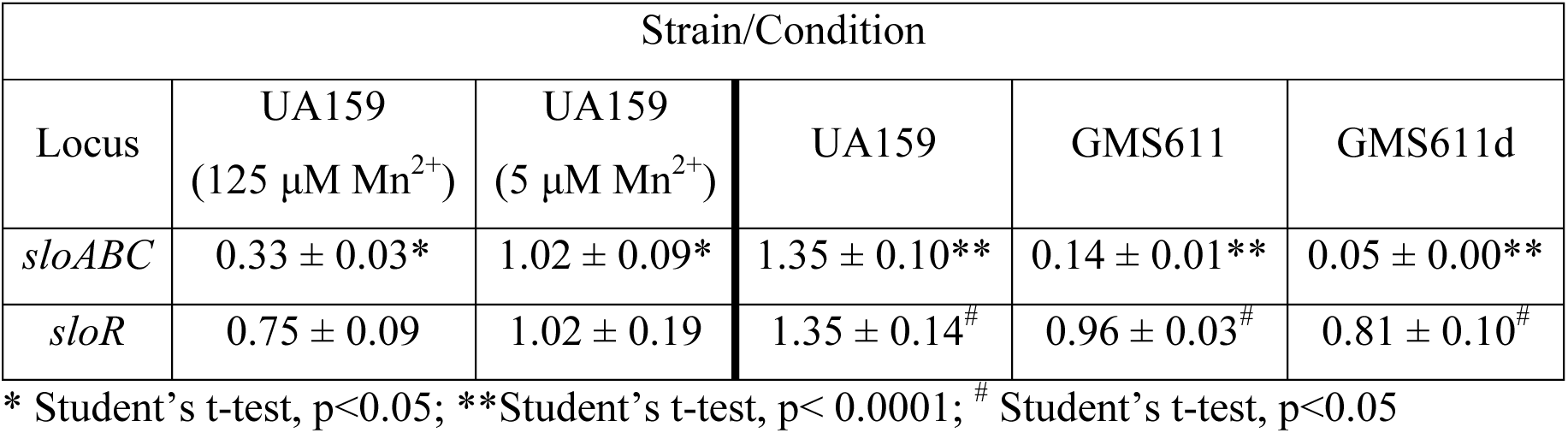
Fold change in expression for *S. mutans sloABCR*.

### The results of 5’ RACE reveal the location of a transcription start site in the intergenic region between the *S. mutans sloC* and *sloR* genes

To investigate whether a promoter might exist on the IGR that separates the *sloABC* and *sloR* genes, we performed 5′ RACE experiments to identify a putative +1 transcription start site. The results revealed that transcription of the 654bp *sloR*-specific transcript begins at an adenosine residue located 19bp upstream of the ATG translation start codon. Mapping the cDNA sequence back to the *S. mutans* UA159 reference genome allowed us to predict and annotate the −10 and −35 sites of the predicted *sloR* promoter (Figure 2). The putative −10 site aligns precisely with the conserved prokaryotic consensus sequence (TATAAT) whereas there is variation in the sequence between the predicted −35 site and its consensus sequence in other prokaryotes. A putative extended −10 element which is characterized by a TGN sequence and the presence of two poly T tracts in the spacer region may compensate for degeneracy in the −35 promoter.

**Figure 2.**
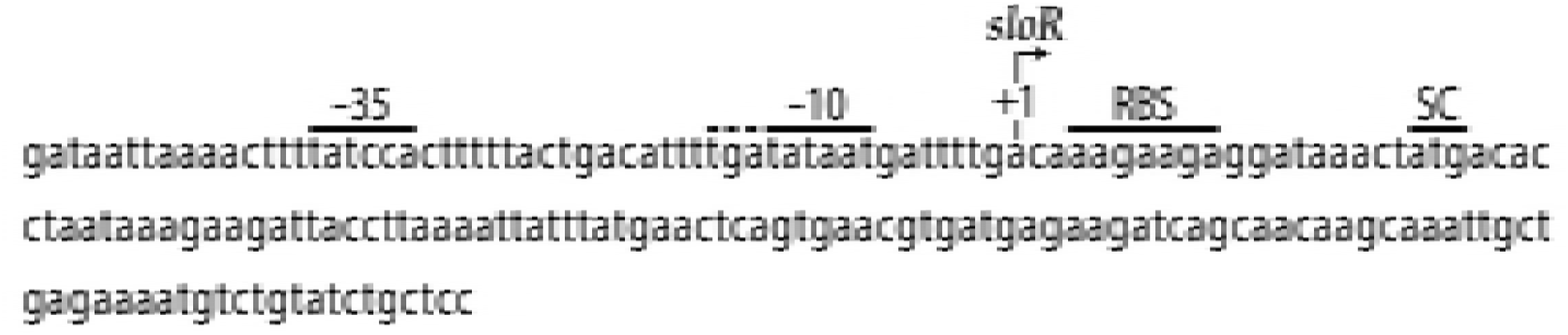
Shown is the intergenic region between the *sloR* and *sloC* genes which harbors a recognizable promoter. The nucleotide sequence of this region was aligned with the *S. mutans* UA159 genome from the NCBI GenBank Database (RefSeq accession number NC_004350.2). The +1 transcription start site (designated by the arrow) marks the transcription start site of the *sloR* gene as defined by 5’RACE, and defines a 19bp 5’ untranslated region (UTR). Also shown are the predicted −35 and −10 promoter regions, the predicted ribosome binding site (RBS), and the start codon (SC) of the 654bp *sloR* gene. A putative extended −10 element is denoted by the dashed line.

### The *S. mutans sloR* gene is transcribed even in the absence of a functional *sloABC* promoter

To investigate the impact of promoter/SRE variants on transcription of the *S. mutans sloABCR* operon, we introduced transition mutations into the 72bp SRE at positions 11 and 11d, both of which share overlap with the −35 and −10 promoter sites upstream of *sloABC*, respectively. Notably, T-to-C mutations at these sites in the resulting GMS611 and GMS611d strain variants culminated in significantly compromised *sloABC* transcription (Student’s t-test, p<0.0001) in qRT-PCR experiments, with Cq values approaching those of the no template and reverse transcriptase-minus controls (data not shown). This is consistent with disruption of the *sloABC* promoter in the GMS611 and GMS611d strain variants. While *sloABC* transcription in these mutant variants was greatly reduced, transcription of *sloR* was diminished to a much lesser extent (Student’s t-test, p<0.05) that we propose is the result of continued transcription from an independent, *sloR*-specific promoter (Table 2).

### The *S. mutans sloR* gene is transcribed from the *sloABC* promoter as well as from an independent promoter on the 184bp IGR

To determine whether *sloR* transcription is indeed driven by an independent *sloR*-specific promoter, we performed reverse transcriptase PCR (RT-PCR) experiments with cDNAs that we generated from the *S. mutans* GMS611 and GMS611d *sloABC* promoter variants and their UA159 wild-type progenitor. As noted above, expression of the *sloABC* operon in the GMS611 and GMS611d is virtually nil, indicating successful disruption of the *sloABC* promoter in these strains. In *S. mutans*, expression of the *sloABC* and *sloR* genes derives from a 3.4Kb polycistronic mRNA owing to transcription off of the UA159 chromosome that is driven by the upstream *sloABC* promoter and transcriptional readthrough of a weak terminator at the 3’ end of the *sloC* gene (21, 32, 33). Herein, we expect to generate a 2.7Kb polycistronic mRNA by RT-PCR given the positioning of the P1 and P3 primers that span the *sloABC* operon and the downstream IGR (Figure 3a). In fact, the results of RT-PCR indicate the presence of a 2.7kb cDNA product in *S. mutans* UA159, and the absence of this product in GMS611 and GMS611d (Figure 3b). In addition to this polycistronic mRNA however, we noted the presence of a 250bp cDNA product in all three *S. mutans* strains with the P2/P3 primer pair. The presence of this amplicon in GMS611 and GMS611d indicates the production of a *sloR*-specific transcript even in the absence of a functional *sloABC* promoter and supports *sloR* transcription from an independent promoter. Importantly, PCR products deriving from genomic DNA when used as the amplification template confirm the specificity of the respective primer pairs (Figure 3b).

**Figure 3.**
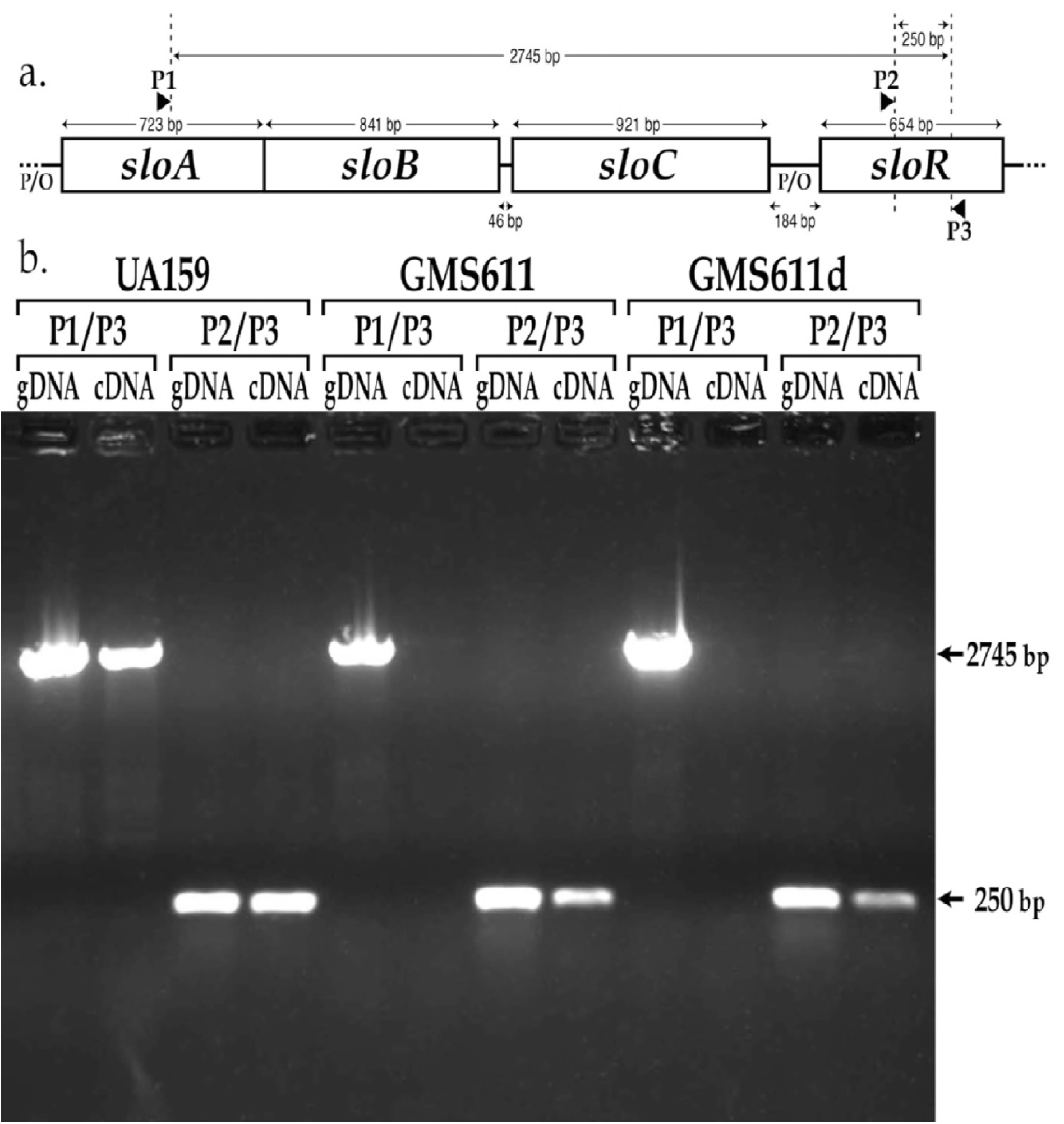
The *S. mutans sloR* gene is transcribed even in the absence of a functional *slo*ABC promoter. **a.)** Map of the *sloABCR* operon and location of primer annealing sites. Primer P1 (*sloA*.RT_PCR.F) anneals within the *sloA* coding sequence, and primers P2 and P3 (*sloR*.RT_PCR.F and *sloR*.RT_PCR.R, respectively) anneal within the *sloR* coding sequence. **b.)** Products of reverse transcriptase PCR resolved on a 0.8% agarose gel. Amplification of cDNA with the P1/P3 primer pair generated a 2745bp amplicon in UA159 but not in GMS611 or GMS611d, consistent with disruption of the *sloABC* promoter in the mutant strains. In contrast, cDNA amplification with the P2/P3 primer pair gave rise to a 250bp amplicon even in the *sloABC* promoter mutants GMS611 and GMS611d, indicating the presence of a *sloR*-specific promoter in the 184bp intergenic region that separates the *sloABC* operon from the downstream *sloR* gene (gDNA = genomic DNA; cDNA = copy DNA).

### SloR binds directly to the intergenic region between the *S. mutans sloC* and *sloR* genes

To determine whether the impact of SloR binding on *sloR* transcription is direct, we performed EMSA experiments, the results of which support direct SloR binding to the intergenic region between the *S. mutans sloC* and *sloR* genes. Specifically, we observed protein-IGR binding when SloR was provided at concentrations as low as 400nM, but not at concentrations of 200nM or less (Figure 4a). This contrasts with the SloR binding we observed at the *sloABC* promoter region which occurred with as little as 60nM SloR protein. These findings support SloR binding to the IGR with lower affinity than that of SloR binding to the *sloABC* promoter region. In addition, SloR-IGR binding was abrogated upon the addition of 1.5mM EDTA, consistent with the metal ion-dependence of the interaction.

**Figure 4.**
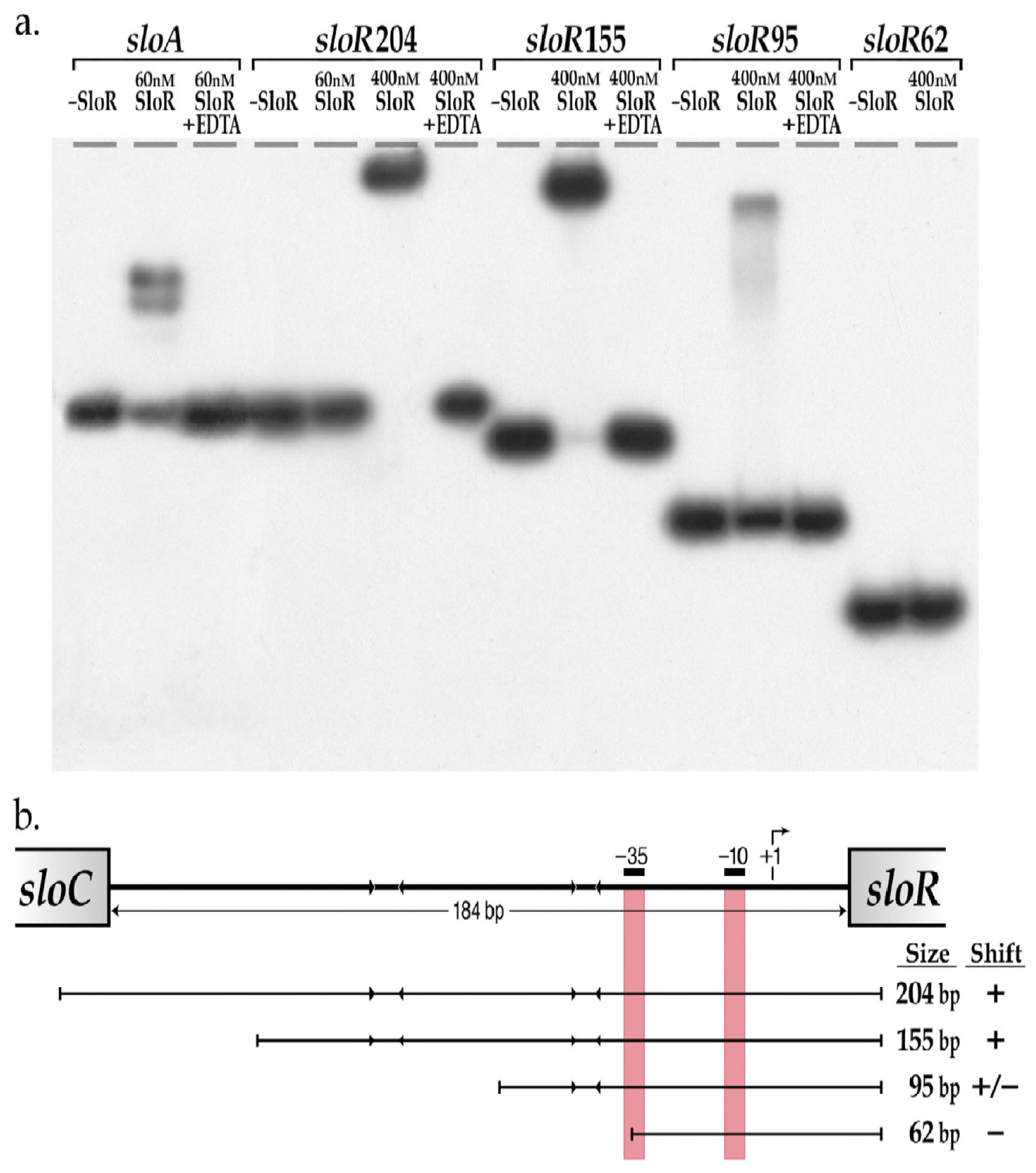
SloR binds directly to the intergenic region (IGR) between the *S. mutans sloC* and *sloR* genes. **a.)** Shown are the results of EMSA which support direct SloR binding to 204bp and 155bp fragments of the *sloC*-*sloR* intergenic region at protein concentrations as low as 400nM. SloR binding to a 95bp IGR derivative was relatively compromised however, and completely absent when a 62bp deletion derivative was used as the binding template. A 205bp amplicon that includes the *sloABC* promoter was used as a positive control for SloR binding. EDTA was added to select reaction mixtures in an attempt to abrogate metal ion-dependent binding. 12% nondenaturing polyacrylamide gels were run at 300 volts for 1.5 hours. Film exposure in the presence of an intensifying screen proceeded for 48 hours at −80°C before development. **b.)** SloR binding to serial deletion fragments of the *S. mutans* IGR. The arrowheads facing inward represent AATTAA hexameric repeats to which SloR putatively binds. The vertical red bars denote the positioning of the −10 and −35 promoter sequences of the s/oR-specific promoter. Whether or not SloR binds to the IGR fragment is shown with a (+) or (−) designation.

We also used EMSA to determine the region on the IGR to which SloR binds. To this end, we generated a series of IGR fragments with serial deletions at their 5’ or 3’ ends (Figure 4a). Interestingly, robust band shifts were generated with DNA fragments harboring promoter-distal hexameric repeat sequences that are located at least 62bp upstream of the +1 transcription start site, but not with DNA fragments less than 62bp from the +1 start site that lack these repeat sequences (Figure 4b).

### Expression of the *S. mutans sloR* gene is subject to positive autoregulation

Direct binding of SloR to promoter-proximal sequences at the *sloABC* locus and to promoter-distal sequences at the *sloR* locus, coupled with their negative and positive effects on global gene expression respectively, led us to hypothesize a role for SloR as a bifunctional regulator in *S. mutans.* To validate such a dual role for SloR, we performed *in vitro* transcription (IVT) experiments, the results of which demonstrate unequivocally that *sloABC* transcription is repressed by SloR while transcription of the *sloR* gene is facilitated by SloR (Figure 5). These findings, quantified with ImageJ software and in combination with the results of binding studies, indicate that SloR can either repress or encourage gene expression via direct binding to DNA. To our knowledge, this is the first experimental evidence to demonstrate bifunctional regulation of gene expression by the SloR metalloregulator in *S. mutans*.

**Figure 5.**
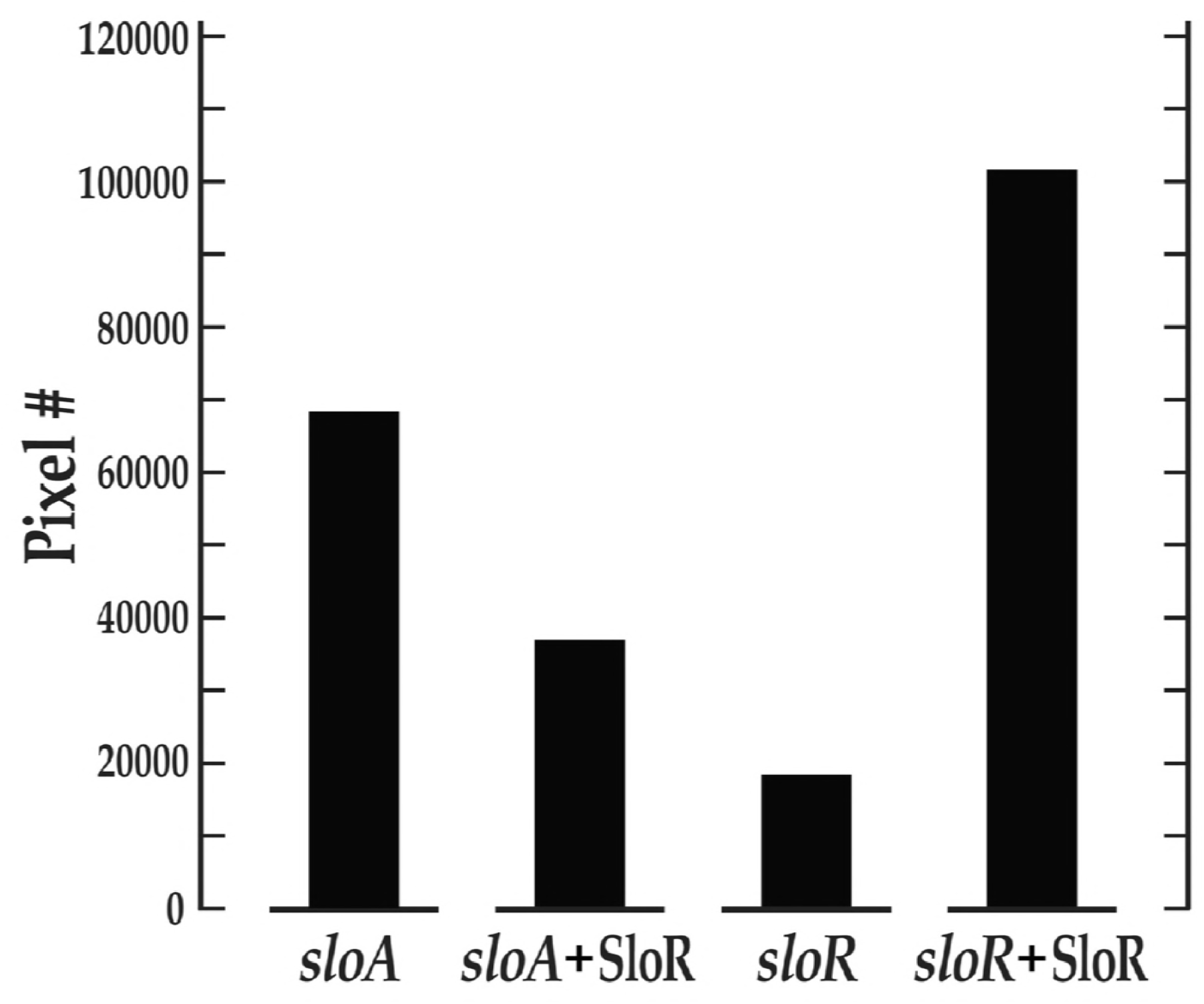
The results of *sloABCR* transcription experiments performed *in vitro* (IVT) support SloR as a bifunctional regulator. Transcription of the *sloABC* and *sloR* genes was quantified using ImageJ software. SloR (75nM) was added to select reaction mixtures containing RNAse inhibitor and either *sloA* or *sloR* template DNA. Pixel counting was performed with ImageJ software and does not include unincorporated α^32^P-UTP. Higher pixel counts indicate greater band intensity. Shown are the results of a single representative experiment (from a total of 3 independent experiments) which support repression of *sloA* and activation of *sloR* transcription by the SloR metalloregulator.

## Discussion

Until recently, work in our laboratory was focused primarily on understanding the mechanism(s) of SloR binding at the *sloABC* locus, which is subject to transcriptional repression by SloR. Herein we present evidence that supports an updated model for SloR binding at this locus, consistent with the direct binding of SloR homodimers to binding sites that span a 72bp region of DNA which includes the −10 and −35 *sloABC* promoter. In previous work (22), we described a pattern for SloR binding at the *sloABC* promoter site that involved two SloR dimers binding to two sets of inverted repeats, each 6bp in length and separated by 8bp. Subsequent analysis of the 72bp region upstream of the *sloABC* operon suggested that, in fact, SloR binds to three distinct but abutting 22bp sites in that region, referred to herein as A, B, and C. Each binding site is comprised of two inverted hexameric repeats with an AATTAA consensus separated by 6bp, thereby defining a putative 6-6-6 motif for SloR binding (Figure 1). This sequence pattern (xxAATTAAxxxxxxTTAATTxx, where “x” is a non-conserved nucleotide) is similar to that of the SloR homolog in *S. gordonii*, called ScaR, which binds to two adjacent 22bp sites upstream of the *scaABC* operon (34) and for which the binding pattern was confirmed by negative staining and electron microscopy (unpublished observations). We therefore expanded what we previously thought to be a 42bp SRE in the *sloABC* promoter region to include at least 30 additional base pairs.

While SloR binds to the central 22bp SRE (region B) with strong affinity when provided as a template in isolation, we measured cooperative interactions between SloR homodimers when bound to adjacent SRE sequences (regions A and C). These cooperative interactions are strong enough to support high affinity binding between SloR homodimers, presumably because of interactions involving the initial binding of SloR homodimers to the B site. Such cooperativity has likewise been observed for the *S. gordonii* ScaR protein (32), suggesting that this property may be a common feature among interactions involving other streptococcal manganese-dependent regulators and their promoter/operator sequences.

In the present study, we extend our SloR binding observations to include the details of protein binding to the IGR that immediately precedes the *S. mutans sloR* gene. Based on accumulating evidence presented herein, we propose that the location of the SloR-DNA binding element relative to the promoter sequences that modulate downstream *sloABC* and *sloR* gene transcription contributes to SloR’s ability to differentially down-regulate *sloABC* promoter activity and up-regulate *sloR* promoter activity. Interestingly, an *in silico* analysis of the 184bp IGR that precedes the *S. mutans sloR* gene failed to reveal a recognizable SRE like the one we describe above for the *sloABC* locus, consistent with differential regulation by SloR at these two loci.

The transcription start site for the *S. mutans sloR* gene occurs within the 184bp IGR that separates *sloR* from the *sloABC* operon immediately upstream, as determined in 5’RACE experiments. From the +1 transcription start site, −10 and −35 promoter regions were predicted and a 19bp 5’ untranslated region (UTR) was defined. Notably, the hexameric −10 region shares 100% sequence identity with the canonical prokaryotic consensus sequence for a −10 promoter (TATAAT) (35, 36). An *in silico* analysis of the *sloR* promoter revealed a putative extended −10 element in the IGR which is absent from the *sloABC* promoter region. Reports in the literature describe such a TGN motif immediately upstream of the −10 sequence as an element that could facilitate downstream transcription by stabilizing the open complex during initiation, and by shortening the distance between the −10 and −35 sites (34, 37). The contact that RNA polymerase makes with the nonamer that defines the extended −10 site could compensate for the suboptimal contact that the polymerase makes with the degenerate −35 sequence (37). The TGN motif that we noted on the *sloR*-containing IGR is similarly located 14-16 nucleotides upstream of the transcription start site. Previous studies have also noted that short poly(T) tracts are characteristic to the spacer region of *E. coli* TG promoters (38). We similarly noted the presence of two poly(T) tracts centered at positions −18 and −29 in the spacer region of *sloR*’*s* TG promoter.

In contrast to the −10 promoter region, the predicted −35 site (TATCCA) shares only 50% sequence identity with the typical prokaryotic promoter sequence (TTGACA) (35). This is not surprising given frequent reports of sequence variation in and around the −35 promoter region within and across prokaryotic species (39). Since promoter strength is, in part, determined by conservation of the −10 and −35 promoter sequences (36) one might expect RNAP to have only moderate binding affinity for the relatively divergent *sloR* promoter (TATAAT and TATCCA) as compared to RNAP binding at the more highly conserved *sloABC* promoter (TATATT and TTGACT) (22), and accordingly, weaker transcription from the former as compared to the latter. The results of DNA binding and expression profiling experiments reported herein support these predictions and suggest a role for divergent *sloABC* and *sloR* promoter sequences in fine-tuning metal ion transport and minimizing the toxic effects of metal ion hyper-accumulation. Additional layers of gene control involving SloR likely evolved at these loci given the importance of maintaining metal ion homeostasis under conditions as transient as those in the oral cavity.

The absence of a recognizable SloR binding motif in the IGR that precedes *sloR* is consistent with differential gene regulation and SloR binding at this locus versus that at the *sloABC* locus. That is, three adjacent palindromes on the 72bp SRE appear to be absent from the IGR, although a pair of consensus palindromes with the sequence AATTAA appear to be uniquely located 44-50bp and 94 −100bp distal to the *sloR* promoter, respectively. Interestingly, reports in the literature describe AT-rich sequences, including palindromes like those in the *sloABC* and *sloR* promoters, that can engender intrinsic curvatures in the DNA (24, 38). To assess inherent DNA curvature in the *sloABC* and *sloR* promoter regions, we applied a BEND algorithm (40) to the 72bp *sloABC* SRE and the 184bp IGR, the results of which revealed high fidelity alignment of AT-rich palindromes with predicted sites for SloR binding (data not shown). Hence, DNA curvature that localizes to the paired palindromic repeats at the *sloABC* and *sloR* loci supports a SloR-DNA interaction that is not strictly defined by nucleotide sequence specificity, but by DNA conformation as well. EMSA studies are currently underway to determine what impact, if any, these AT-rich palindromes may have on SloR-DNA binding, and whether the DNA curvature they can instigate contributes to SloR’s function as a repressor versus an enhancer of gene transcription.

Differential expression of the *sloABC* and *sloR* genes was especially pronounced in *in vitro* transcription (IVT) assays where we used the 5’ end of the *sloA* or *sloR* coding regions and up to 200bp of upstream DNA sequence as the DNA template. Pixel counting of the resulting mRNA transcripts on autoradiograms, performed with Image J, revealed considerably more robust *sloR* transcription in the presence of exogenous SloR than in its absence (Figure 5). This contrasts with the *in vitro* transcription we observed for the upstream *sloABC* operon which, as expected, was repressed by the presence of SloR. Taken together with the EMSA results that support direct SloR binding at these loci, these data demonstrably support a role for SloR as a bifunctional regulator of *S. mutans* gene transcription. While the mechanism for *sloABC* repression likely derives from SloR binding to an SRE that shares overlap with the *sloABC* promoter, thereby excluding RNAP from promoter access, *sloR* transcription is the likely result of de-repression with SloR binding to promoter-distal sites. In future work, we will consider Mn^2+^ status as a potential contributor to differential SloR binding and gene transcription outcomes. In fact, binding of the MntR metalloregulator to different sequences upstream of the *mntABCD* locus in *Bacillus subtilis* is known to be Mn^2+^ concentration-dependent (41).

In summary, the results of the present study support SloR-mediated transcriptional events at the *sloR* locus that are different from those at the *sloABC* locus and lend further support to a role for SloR as a bifunctional regulator of gene transcription. It’s tempting to suggest that the *sloR*-specific promoter on the 184bp IGR evolved to ensure at least some level of constitutive SloR production. Accordingly, when free metal ions are introduced into the oral cavity during a mealtime, *S. mutans* is poised and ready to modulate the controlled uptake of the exogenous Mn^2+^ and Fe^2+^ it needs for survival. Hence, while the sudden introduction of metal ions into the mouth could prove damaging to some constituents of the oral microbiota, *S. mutans* can exploit these conditions with a metal ion-dependent SloR regulator that can repress the cytotoxic effects of excessive metal ion import, while maintaining baseline levels of SloR from an ancillary promoter. Such fine-tuned gene regulation can impact cell function beyond the scope of metal ion homeostasis, and influence processes like adherence, acid production and the oxidative stress response that more directly contribute to *S. mutans*-induced disease (7, 8, 24, 25). In conclusion, we propose that *S. mutans* coordinates the regulated expression of its metal ion transport machinery with that of its virulence attributes. An improved understanding of *sloR* autoregulation is significant because it can elucidate the mechanisms that fine-tune the regulated expression of metal ion homeostasis and virulence in an important oral pathogen. Moreover, from these investigations we may better elucidate the details of SloR-mediated gene regulation that can benefit the design of an anti-SloR therapeutic aimed at alleviating and/or preventing caries.

## Materials & Methods

### Bacterial strains, plasmids, and primers

The bacterial strains used in this study are listed in Table 1. Working stocks of each bacterial strain were prepared from overnight cultures and stored in 20% or 50% sterile glycerol at −20°C or −80°C, respectively.

The primers used in this study are shown in Table 1. All primers were designed using the Primer Blast tool from the NCBI website. A RefSeq record was used as the input with forward and reverse primer locations specified by the user. Lack of secondary structure was confirmed using the Oligo Evaluator tool (Sigma) and primers were checked for specificity against the *S. mutans* UA159 genome with the NCBI Basic Local Alignment Search Tool (BLAST).

### Negative-staining and EM analysis

A SloR-Mn^2+^-72bp SRE complex was prepared *in vitro* assuming a 3:1 stoichiometry of dimers to DNA. The reaction mixtures were then systematically diluted to 25, 50 and 100nM for negative-stain EM grid preparation using 1% uranyl acetate. After screening for the optimal staining quality and particle concentration, single-particle data were collected on the 50 nM specimen grid using an FEI T12 electron microscope at 120KeV and 68,000x nominal magnification, producing 108 micrographs at 3.15 Å/pixel in the image with varying defocus between 0.9 and 1.8 μm. Then, 8,500 particles of the SloR-DNA complex were selected for 2D classification using the PARTICLE (www.sbgrid.org/software/titles/particle) program.

### Construction of *S. mutans* promoter variants

To generate specific mutations in the promoters that drive *sloABC* transcription, we adopted a markerless mutagenesis strategy similar to that described by Xie *et al.* (27). This involved constructing promoter variants with an IFDC2 cassette inserted within the *sloABC* promoter region for subsequent CSP-transformation into *S. mutans* UA159 to generate the erythromycin-resistant and p-4-chlorophenylalanine-sensitive GMS602 strain. The double-crossover event was confirmed by polymerase chain reaction (PCR) and Sanger sequencing. Derivatives of GMS602 were generated by overlap extension PCR (OE-PCR) with the reverse primers harboring a single point mutation in the predicted 72bp SRE that precedes the *sloABC* genes (Table 1). *S. mutans* GMS611 was generated with degenerate primers that introduced a point mutation into the SRE at nucleotide position 11 that we predict shares overlap with the *sloABC*-specific −35 promoter region (22). Likewise, GMS611d was generated with a different degenerate primer set that introduced a point mutation at position 11d into the SRE, which we predict shares overlap with the *sloABC*-specific −10 promoter region. Thymine to cytosine transition mutations were generated in both *S. mutans* strains and validated by Sanger sequencing (22).

### Chromosomal DNA Isolation

*S. mutans* grown overnight at 37°C and 5% CO_2_ in 14ml Todd-Hewitt Yeast Extract (THYE) broth were pelleted by centrifugation at 7000rpm for 5 minutes in a Sorvall RCB centrifuge after which the cells were resuspended in 1ml Tris-EDTA buffer (10mM Tris, 1mM EDTA) and chemically disrupted according to established protocols (8, 22). Genomic DNA was purified in subsequent rounds of phenol-chloroform extraction, ethanol precipitated, and resuspended overnight in nuclease-free water (Ambion) at 4°C with gentle agitation (28). Nucleic acid yield and purity were assessed with a NanoDrop Lite spectrophotometer (Thermo Fisher Scientific) and the samples were stored at −20°C.

### RNA Isolation

RNA was isolated from *S. mutans* strains according to established protocols (8). Cells were grown to mid-logarithmic phase (OD_600nm_ = 0.4-0.6) before pelleting by centrifugation as described above and resuspending in RNAProtect (Qiagen). Total intact RNA was purified following cell lysis and DNase I treatment with a Qiagen RNeasy kit after which nucleic acid yield and purity were assessed with a NanoDrop Lite spectrophotometer (Thermo Fisher Scientific). RNA samples were analyzed for integrity by agarose gel electrophoresis and stored at −80°C.

### RT-PCR

Reverse-transcriptase PCR was performed with cDNAs that were reverse-transcribed from RNA using a First Strand cDNA Synthesis kit according to the recommendations of the supplier (Thermo Fisher Scientific) or else with chromosomal DNA isolated from *S. mutans* UA159, GMS611, or GMS611d according to a modification of Idone *et al.* (29). Q5 Hi-Fidelity Polymerase was used for PCR in accordance with the recommendations of the supplier (New England Biolabs). Each 50 μL PCR reaction consisted of 10 μL 5X Q5 Reaction buffer, 1 μL 10mM dNTPs, 2.5 μL *sloA*.RT_PCR.F or *sloR*.RT_PCR.F, 2.5 μL *sloA*.RT_PCR.R or *sloR*.RT_PCR.R [Table 1]), 200 ng of genomic DNA or 1 μL of cDNA product, nuclease-free water up to 49.5 μL, and 0.5 μL of Q5 Hi-Fi Polymerase added to the reaction just prior to amplification. PCR conditions were as follows: initial denaturation at 98°C for 30 s, followed by 25 cycles of denaturation at 98°C for 10 s, annealing at 67°C for 30 s, and extension at 72°C, concluding with a final extension at 72°C for 80 s. An aliquot of each sample was visualized on 0.8% agarose gels as described above.

### 5’ RACE

To reveal the *sloR* transcription start site and predict the −10 and −35 regions of the *sloR* promoter we performed 5′ Rapid Amplification of cDNA Ends (5’ RACE) with an Invitrogen 5′ RACE Kit in accordance with the manufacturer’s protocol unless otherwise specified. RNA isolated from *S. mutans* GMS611 and GMS611d was reverse-transcribed into cDNA using a *sloR*.[R].GSP1.A reverse primer and 200 U of Superscript II Reverse Transcriptase. After S.N.A.P. column purification, the cDNA was poly(-dC) tailed using terminal deoxynucleotidyl transferase as described in the manufacturer’s protocol. The cDNA products were PCR amplified using the kit-supplied Abridged Anchor Primer (AAP), a *sloR*.[R].GSP2.B reverse primer, and Platinum Hi-Fi *Taq* polymerase, in a BioRad PCR machine programmed at 94°C for 2 minutes, followed by 35 cycles of 94°C for 15 s, 58°C for 30 s, and 68°C for 1 minute, and a final hold at 4°C. An aliquot of the resulting PCR product was analyzed by agarose gel electrophoresis and the remainder was purified using a MinElute PCR Purification kit in accordance with the recommendations of the supplier (Qiagen). The purified amplicons were quantified using a NanoDrop Lite spectrophotometer (Thermo Fisher Scientific) and sequenced (Eurofins) using the reverse primers *sloR*.GSP2.A or *sloR*.nested.GSP. The 5′ sequence of the mRNA transcript was aligned with the *S. mutans* UA159 reference genome (RefSeq accession number NC_004350.2) from NCBI to identify the nucleotide that immediately follows the poly(-dC)-tail as the transcription start site (+1 site), predict the −10 and −35 promoter sequences in the 184bp intergenic region, and reveal the 5′ untranslated region of the *sloR* transcript (30).

### EMSA

Electrophoretic mobility shift assays (EMSAs) were performed according to established protocols to determine whether SloR binding to the intergenic region upstream of the *sloR* gene is direct, and to narrow down the region of SloR binding at this locus (8, 22, 23). Primer design for the DNA binding template spanned the 184bp intergenic region (IGR) between *sloC* and *sloR* and included serial deletions thereof. PCR amplification was performed with Q5 polymerase according to the manufacturer’s protocol (New England Biolabs) using the following thermal cycling conditions: initial denaturation at 98°C for 30 s, 35 cycles at 98°C for 10 s, annealing at 60°C for 30 s, and extension at 72°C for 30 s, with a final extension at 72°C for 2 minutes. The resulting amplicons were PCR purified as described earlier, confirmed by agarose gel electrophoresis, and quantified by NanoDrop spectrophotometry.

The resulting amplicons were end-labeled with [γ-^32^P]-dATP (Perkin-Elmer) in the presence of T_4_ polynucleotide kinase (New England BioLabs) after which they were centrifuged through a TE Select-D G-25 spin column (Roche Applied Science) to remove the unincorporated ^32^P-dATP. Binding reactions were prepared as described previously (23) in a 16-μl total volume containing 1 μl of end-labeled amplicon, purified native SloR protein at concentrations ranging from 60 nM to 400nM, and 3.2 μl of 5× binding buffer (42 mM NaH2PO4, 58 mM Na2HPO4, 250 mM NaCl, 25 mM MgCl_2_, 50 μg/ml bovine serum albumin, 1 mg sonicated salmon sperm DNA, 50% [vol/vol] glycerol, and 37.5 μM MnCl_2_). EDTA was added to a separate reaction mixture at a final concentration of 1.5mM to validate SloR binding that is metal ion-dependent. An end-labeled *sloA* promoter-containing amplicon was used as a positive control for SloR binding (7, 22, 23). Samples were loaded onto 12% nondenaturing polyacrylamide gels (3 ml 20× bis-Tris borate [pH 7.4], 74 μl 100 nM MnCl_2_, 1.5 ml 100% glycerol, 24 ml 30% acrylamide [37.5:1 acrylamide-bis], 31 ml Millipore H_2_O, 300 μl 15% ammonium persulfate [APS], 90 μl TEMED [*N*,*N*,*N*′,*N*′-tetramethylethylenediamine]) and resolved at 300 volts for 1.5 hours. Gels were exposed to Kodak BioMax film for 24-72 hours at –80°C in the presence of an intensifying screen prior to autoradiography.

### Fluorescence anisotropy

Equilibrium binding of SloR to regions within the *sloABC* promoter/operator region was probed by titrating SloR onto fluoresceinated DNA and monitoring binding by fluorescence anisotropy. Duplex DNA fragments containing relevant sequences were prepared from oligonucleotides obtained from Integrated DNA Technologies (Coralville, IA). A 5’ fluoresceinated oligonucleotide was annealed with a 10% excess of its unlabeled complementary strand in 25mM HEPES, pH7.9, 50mM NaCl by heating to 90°C and cooling slowly to room temperature (Table S1). Titrations were performed in either high or low salt buffers; the high salt buffer contained 25mM HEPES, pH 8.0, 250mM NaCl, 10% glycerol and 1mM MnCl_2_ whereas the low salt buffer contained 25mM HEPES, pH7.9, 50mM NaCl, and 1mM MnCl_2_. SloR was titrated into 1ml of 1nM fluoresceinated duplex DNA in buffer and anisotropy measurements were made at 25°C using a Beacon 2000 fluorescence polarization instrument. Data were fit to one of several equations related to simple 1:1 binding stoichiometry, where K_d_ was greater than 10nM (equation 1), less than 10nM (equation 2) or to the Hill equation (equation 3).

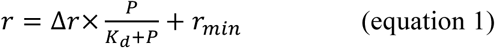

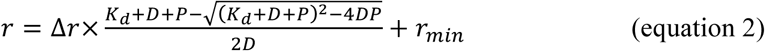

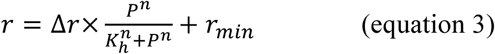

In these equations, r is anisotropy, Δr is the total change in anisotropy at saturation, K_d_ is the dissociation constant, K_h_ is the concentration of SloR dimers giving 50% maximal binding under cooperative conditions, n is the Hill coefficient, and r_min_ is the anisotropy obtained before addition of SloR. P is the concentration of SloR dimers and D is the concentration of duplex DNA. In addition, where non-specific binding prevented signal saturation, a term to model the slow linear increase in anisotropy was added to equations 1 or 2, with the form K_ns_P where K_ns_ is the non-specific binding constant. All curve fitting was performed in R software.

### Preparation of *S. mutans* RpoD

An *E. coli* DH5α strain harboring plasmid pIB611 (kind gift of Indranil Biswas, University of Kansas) was streaked for isolation on L-agar plates supplemented with ampicillin (100 μg/mL) and incubated overnight at 37°C. Resident on pIB611 is the *S. mutans rpoD* gene cloned directly downstream of an inducible operon on vector pET-23d(+) and upstream of a C-terminal His-tag (31). Importantly, the 6x His-tag was shown in *in vitro* transcription experiments to have no physiological impact on RpoD functionality (31). Plasmid pIB611 was purified with a Qiagen miniprep kit according to the recommendations of the supplier and mapped by restriction digestion (New England Biolabs).

Next, pIB611 was used to transform *E. coli* BL21 DE3 cells in accordance with the manufacturer’s protocol (Invitrogen). Successful transformants were selected after overnight growth on L-agar plates supplemented with 100ug/mL ampicillin and used to inoculate a 25 mL starter culture for yet another overnight incubation. This culture was then used to inoculate 500 mL of pre-warmed L-ampicillin (100ug/ml) broth in a 2.8 L Fernbach flask which was incubated at 37°C with continuous shaking at 200rpm. When the cells reached mid-log phase (OD_600_=0.4), isopropyl β-D-1-thiogalactopyranoside (IPTG) was added at a final concentration of 0.5mM to induce expression of the T7 RNA polymerase in the BL21 DE3 cells. Protein induction proceeded with continuous shaking for an additional 3.5 hours after which the cells were centrifuged as previously described and stored as dry pellets at −80°C.

Cell pellets were resuspended in His-Binding Buffer (0.5M NaCl, 20mM Tris-HCl, 5mM imidazole) with Halt EDTA-free protease inhibitor at 1x concentration (Thermo Fisher Scientific). The cell suspension was sonicated (Ultrasonic Power Corporation, model number: 2000U) at 60% power for six 30-second cycles using a 0.5 second pulse, with samples maintained on ice between runs. Cells were pelleted by centrifugation at 10,000 x *g* for 30 minutes at 4°C, after which aliquots of the supernatant and 2X Laemmli buffer (4% SDS, 20% glycerol, 10% 2-mercaptoethanol, 0.004% bromophenol blue and 0.125 M Tris HCl, pH 6.8) were mixed in equal proportions and resolved on 10% Bis-Tris polyacrylamide gels in 3-(N-morpholino) propanesulfonic acid (MOPS) buffer. Proteins were visualized with Sypro Ruby Gel Stain (Thermo Fisher Scientific) according to the manufacturer’s instructions. Polyacrylamide gels were fixed for 45 minutes in fixative solution (50% methanol, 7% acetic acid) with gentle shaking on an orbital shaker. The fixative was subsequently decanted and replaced with 80 mL of Sypro Ruby Gel Stain. Gels were covered and left to stain on an orbital shaker overnight at room temperature. The gel stain was decanted and wash solution (10% methanol, 7% acetic acid) added before UV visualization.

The remaining cell lysate was further purified by Ni^2+^-Nitrilotriacetic acid (Ni-NTA) column chromatography (Thermo Fisher Scientific) at 4°C according to the manufacturer’s instructions. The columns were placed on a rotating platform for 30 minutes at 4°C to encourage RpoD binding to the Ni-NTA resin before centrifugation to remove unbound protein from the column. After elution, protein yield was determined using both NanoDrop Lite spectrophotometry and a bicinchoninic acid (BCA) protein determination assay (Thermo Fisher Scientific). RpoD purity was assessed on SDS-PAGE gels following Sypro Ruby staining. Select fractions containing RpoD were dialyzed using G2 Slide-A-Lyzer Cassettes with a 10 kDa molecular weight cut off (Thermo Fisher Scientific) in dialysis buffer (25% glycerol in 1x PBS). RpoD was assayed for concentration as described above and stored at −20°C.

### *In vitro* transcription

*In vitro* transcription was performed according to an adaptation of Kajfasz et al (9). First, genomic DNA spanning approximately 100-200bp of the *sloA* or *sloR* coding region and about 150-200bp of upstream sequence was amplified by PCR with Q5 polymerase (New England Biolabs) and either PM.IVT.*sloA*.F and PM.IVT.*sloA*.R or PM.IVT.*sloR*.F and PM.IVT.*sloR*.R (Table S1) according to the following thermal cycling conditions: initial denaturation at 98°C for 30 s, 35 cycles at 98°C for 10 s, annealing (60°C for *sloA*, 67°C for *sloR*) for 30 s, and extension at 72°C for 30 s, with a final extension at 72°C for 2 minutes. The resulting amplicons were PCR purified as described earlier and confirmed by agarose gel electrophoresis. DNA concentration was determined spectrophotometrically on a NanoDrop Lite spectrophotometer. Next, in 1.5 mL microfuge tubes, 10 nM DNA template (*sloA* or *sloR*), 1U *E. coli* RNA Polymerase core enzyme (New England Biolabs), 20U of SUPERase RNase Inhibitor (Thermo Fisher Scientific), 25 nM *S. mutans* RpoD extract (based on BCA assay), +/− 75nM purified SloR were mixed in Reaction Buffer (10 Mm Tris-HCl [pH 8.0], 50 mM NaCl, 5 mM MgCl_2_, 50 μg/mL bovine serum albumin) to yield a final volume of 17.8 μL and incubated at 37°C for 10 minutes. To generate an mRNA transcript, 2.2 μL of Nucleotide Mixture (200 μM ATP, 200 μM GTP, 200 μM CTP, 10 μM UTP) and 5 μCi α^32^P-UTP (Perkin Elmer) were then added and the reaction was incubated at 37°C for 10 minutes. 10 μL of Stop Solution (1M ammonium acetate, 0.1 mg/mL yeast tRNA (Ambion), 0.03M EDTA) was added to terminate transcription, after which 90 μL of ice-cold 99% EtOH was added for ethanol precipitation overnight at 4°C. The following day samples were pelleted at 16,200 x *g* for 30 minutes, followed by three rounds of washing with 70% EtOH, with additional centrifugation between washes. A final wash with 99% EtOH was performed after which the samples were lyophilized in a vacuum centrifuge (Eppendorf) for 10 minutes. The radiolabeled cell pellet was resuspended in 5 μL of formamide dye (0.3% xylene cyanol, 0.3% bromophenol blue, 12mM EDTA, dissolved in formamide) and the samples were heated to 70°C in a water bath for 2 minutes before placing on ice for gel loading. Samples were resolved on a Novex 10% TBE-7M Urea Gel in 1X TBE Buffer (10.8 g Tris, 5.5g boric acid, 0.01M EDTA [pH 8.0]) and mRNA transcripts were visualized via autoradiography using BIOMAX XAR Film (Thermo Fisher Scientific) exposed for up to 4 hours at −80°C in the presence of an intensifying screen. Film was developed according to standard protocols and ImageJ software was used to quantify the band intensities between samples.

### Semi-quantitative real-time PCR (qRT-PCR)

Total intact RNA was isolated from mid-logarithmic phase *S. mutans* cultures of the wild-type UA159 strain grown in a semi-defined medium (SDM) (22) supplemented with either 5uM (low) or 125uM (high) MnSO_4_. The resulting RNAs were analyzed for integrity on 0.8% agarose gels before reverse-transcribing 100ng of each RNA sample into cDNA as described above. The cDNAs were used as templates for qRT-PCR which was performed in accordance with established protocols (22) in a CXR thermal cycler (BioRad). Specifically, *sloABC* and *sloR* transcription was assessed in three independent qRT-PCR experiments, each performed in triplicate and normalized against the expression of *gyrA* (8,22).

To assess the impact of transition mutations in the SRE that precedes the *sloABC* operon on downstream *sloR* transcription, *S. mutans* GMS611 and GMS611d were grown as described above except without supplemental Mn^2+^. Total intact RNA was isolated and reverse transcribed as described, and the results of qRT-PCR were normalized against *hk11*, the expression of which does not change under the experimental test conditions.

## Acknowledgements

This research was supported by NIH Grant #DE014711 to G.A.S., the T. Ragan Ryan Summer Research Fund to P.M., and by the Middlebury College Department of Biology. We acknowledge Mr. Gary Nelson for figure preparation and Dr. Indranil Biswas for providing plasmid pIB611. Many thanks to Dr. Robert Haney for his bioinformatics expertise and assistance with the microarray data, to Dr. Jessica Kajfasz for her guidance with the IVT experiments, and to Mr. Frank Spatafora for general technical assistance. We declare no conflicts of interest for this work.

